# Diurnal changes in glutamate/glutamine levels of healthy young adults assessed by proton magnetic resonance spectroscopy

**DOI:** 10.1101/216762

**Authors:** C. Volk, V. Jaramillo, RR. Merki, RL. O’Gorman, R. Huber

## Abstract

Molecular and electrophysiological studies suggest that sleep ensures efficient functioning of the brain by maintaining synaptic homeostasis. The glutamate receptor α-amino-3-hydroxy-5-methyl-4-isoxazolepropionic acid (AMPA receptor) is involved in synaptic plasticity processes and it was shown that its expression changes across the sleep wake cycle. Moreover, animal studies have shown that glutamate levels are reduced during non-rapid eye movement (NREM) sleep and that the rate of the decrease is positively correlated with sleep EEG slow wave activity (SWA). In this study, we aimed to assess if proton magnetic resonance spectroscopy (^1^H-MRS) is sensitive to diurnal changes of glutamate + glutamine (GLX) in healthy young adults and if potential overnight changes of GLX are correlated to SWA. ^1^H-MRS spectra of 16 adult subjects were measured in the parietal lobe in the evening and in the subsequent morning using a 3T MRI scanner. The night between the scans was recorded with high-density EEG. Our results revealed a significant overnight reduction in GLX levels, whereas other metabolites did not show any significant change. Moreover, the decrease in GLX was positively correlated with the decrease of SWA in the course of the night. Our study demonstrates that quantification of diurnal changes in GLX is possible by means of ^1^H-MRS and indicates a relationship between changes in GLX and SWA, a marker that is closely linked to the restorative function of sleep. This relationship might be of particular interest in clinical populations in which sleep is disturbed.

## Introduction

It is well known that sleep is essential for healthy brain function and even a moderate lack of sleep can interfere with basic neurobehavioral functions (Krause et al., 2017, Van Dongen et al., 2003). Although the exact mechanism underlying the restorative nature of sleep is still under debate, insights from molecular and electrophysiological studies suggest that sleep ensures efficient functioning of the brain by maintaining neuronal homeostasis. During wakefulness, we constantly interact with the environment, which leads to a progressive build-up of synaptic strength (Tononi and Cirelli, 2014). A continuous increase in synaptic strength would lead to 1) a saturation in neural plasticity, limiting the brain’s capacity to process new inputs (Turrigiano, 2008), 2) increase energy demands to an unsustainable level, necessitating increased cellular maintenance (Vyazovskiy and Harris, 2013) and increased removal of metabolic waste products (Xie et al., 2013). The mechanism thought to underlie the down-regulation of synaptic strength has been linked to deep sleep. During non-rapid eye movement (NREM) sleep, when large populations of cortical neurons oscillate synchronously between a phase of depolarisation (on-state) and hyperpolarisation (off-state), this oscillation becomes visible in the surface EEG in form of sleep slow waves (oscillations between 0.5 and 4.5 Hz; Vyazovskiy et al., 2009). Slow wave activity (SWA, EEG power between 1 - 4.5 Hz) increases with the time spent awake and decreases in the course of the night (Borbély and Achermann, 2005). Additionally, SWA was shown to increase in a use-dependent manner: Cortical areas that have actively been used during the day (by practicing specific tasks) show higher SWA in the following night (Hanlon et al., 2009, Huber et al., 2004) and areas deprived of peripheral input display a decrease in SWA (Huber et al., 2006). Thus, SWA is tightly linked to the restorative function of sleep.

A major molecular marker of changes in synaptic strength is the glutamate receptor α-amino-3-hydroxy-5-methyl-4-isoxazolepropionic acid (AMPA) receptor. Changes in the AMPA receptor function represent a key mechanism of synaptic plasticity (Huganir and Nicoll, 2013, Lee and Kirkwood, 2011) and it was shown that AMPA receptor levels in the rat brain are high during wakefulness and low after sleep (Vyazovskiy et al., 2008). Concordantly, an in vivo amperometry study in rats showed that glutamate (GLU) concentrations increase during wakefulness and rapid-eye movement (REM) sleep and decrease during NREM sleep (Dash et al., 2009). Moreover, the decrease of GLU during NREM sleep positively correlated with SWA, suggesting that NREM sleep is essential to keep GLU in a homeostatic range (Dash et al., 2009). The goal of our study was to specifically assess if this relation between GLU levels and SWA is also present and measureable in the human brain. Proton magnetic resonance spectroscopy (^1^H-MRS), a non-invasive method for investigating biochemical changes in the living human brain, is widely used in both clinical and research settings to investigate changes in a variety of metabolites, including neuronal amino acids such as gamma-Aminobutyric acid (GABA), glutamate + glutamine (GLX) or N-acetylaspartate (NAA; Bollmann et al., 2015, Huang et al., 2015, Rowland et al., 2013). In this study, we evaluated whether ^1^H-MRS is sensitive to changes in GLX from evening (after a day of being awake) to morning (after a night of sleep) in healthy young adults. Additionally, we acquired a high-density sleep electroencephalogram (hd sleep EEG) in the night between the two ^1^H-MRS sessions in order to relate potential overnight changes in GLX levels to SWA. We hypothesized that GLX concentrations in the human brain would show an overnight reduction.

## Materials and methods

### Participants

18 adults between 18 and 24 years of age were recruited via advertisements placed at the University. 2 subjects had to be excluded due to low sleep quality (sleep efficiency < 80 %). The remaining 16 subjects (mean ± s.e.m., 21.3 ± 2.2 years, 6 females), fulfilled the following inclusion criteria: No personal or family history of psychopathology, no sleep disorders, good sleepers (sleep efficiency > 80 %), no chronic diseases, no current use of psychoactive agents or other medications, no travelling across the time zone 1 months before the study, no high caffeine (> 160 mg caffeine / day) or alcohol (> 14 mg alcohol / day) consumption, non-smoker. Subjects had to refrain from alcohol starting 48 hours prior to the experiment and extensive exercise and sauna was not allowed on the day of the recordings. The study was approved by the local ethics committee and subjects gave written informed consent before participating. Participants received a monetary compensation.

### Experimental procedure

One week prior to the assessment, participants were instructed to keep a regular sleep-wake schedule according to their habitual bed time. Compliance was verified with self-reported sleep logs and wrist motor actigraphy (GENEActiv, activinsights Ltd., Kimbolton, Huntingdon, UK). On the night of the sleep recording, subjects arrived ~ 3 hours before their usual bedtime at the sleep laboratory of the University Children’s Hospital Zurich. The experimental procedures started with the magnetic resonance imaging (MRI) around 2 hours before the subject’s habitual bedtime. Afterwards, participants were equipped with the hd EEG net and sleep was recorded all night. Next morning, after removal of the EEG net, the MRI was repeated (~ 1 hour after lights on).

### High-density sleep electroencephalography

All-night sleep EEG was recorded with a high-density EEG net (128 channels, Sensor Net for long-term monitoring; Electrical Geodesic Inc., EGI, Eugene, OR, USA). In addition, two submental EMG electrodes for visual scoring and two electrodes at the earlobes were applied (gold, Grass Technologies, West Warwick, RI, USA). The nets were adjusted to the vertex and mastoids and subsequently filled with electrolyte gel. Impedances were kept below 50 kΩ. EEG recordings were sampled at 500 Hz (filtered between .01 and 200 Hz) and referenced to the vertex (Cz). Subsequently, data were band-pass filtered (0.5 - 50 Hz) and down sampled to 128 Hz. Sleep stages were scored for 20-sec epochs according to standard criteria (Iber et al., 2007) by a sleep expert and verified by a second sleep expert. Artefacts in the 20 second epochs were removed by visual inspection and if power exceeded a threshold based on a mean power value in the 0.75 - 4.5 or 20 - 30 Hz band. Channels with bad quality and channels below the ears (due to common contamination by muscle artefacts) were removed and data was re-referenced to the average value of all good quality channels. For further analyses, sleep cycles were defined according to the criteria of Feinberg and Floyd (Feinberg and Floyd, 1979) and spectral analysis of consecutive 20 second epochs for each sleep cycle (fast Fourier transformation, Hanning window, average of five 4 second epochs, frequency resolution of 0.25 Hz) was performed. Missing data from excluded electrodes were interpolated using spherical linear interpolation (Delorme and Makeig, 2004) resulting in 109 channels per subject. SWA was calculated as the mean power in the frequency band between 1 and 4.5 Hz for every NREM sleep episode (stage N2 and stage N3).

### Magnetic resonance imaging

MR imaging and spectroscopy scans were performed with a 3T GE MR 750 scanner, using an 8 channel receive-only head coil. Single voxel Point RESolved (PRESS) ^1^H MR spectra were acquired from a 20 x 20 x 20 mm^3^ voxel of interest positioned in the left parietal lobe, as described previously (Robertson et al., 2001, see figure 1A). The parietal lobe was chosen because it includes a large proportion of grey matter and the voxel can be positioned according to a standard set of measurements from anatomical landmarks, ensuring a reproducible voxel position between participants regardless of head position. Spectra were acquired with an echo time (TE) of 35 ms, a repetition time (TR) of 3 seconds, and 96 spectral averages. The scanning protocol also included a 3D high resolution T1-weighted IR-SPGR scan (TE = 3 ms, TR = 8 ms, inversion time = 600 ms, flip angle = 8 degrees, voxel resolution = 1mm^3^), used for correction for partial volume CSF effects. Water-scaled metabolite concentrations were calculated with LCModel version 6.3-1k, and corrected for partial volume CSF contamination after segmentation of the 3D T1-weighted images into grey matter (GM), white matter (WM), and CSF maps in SPM8 (Statistical Parametric Mapping; Wellcome Department of Imaging Neuroscience, Institute of Neurology, University College London). Each spectrum was visually inspected for the presence of artefacts or fitting errors, and spectra with Cramer-Rao variance bounds of more than 10 % for Creatine or N-acetyl-aspartate, or more than 20 % for glutamate were excluded from further analysis. Due to the tight spectral overlap between glutamate and glutamine at 3T, glutamate levels were quantified as GLX (the sum of glutamate and glutamine), as frequently done (e.g. Dlabac-de Lange et al., 2017).

**Figure 1.**
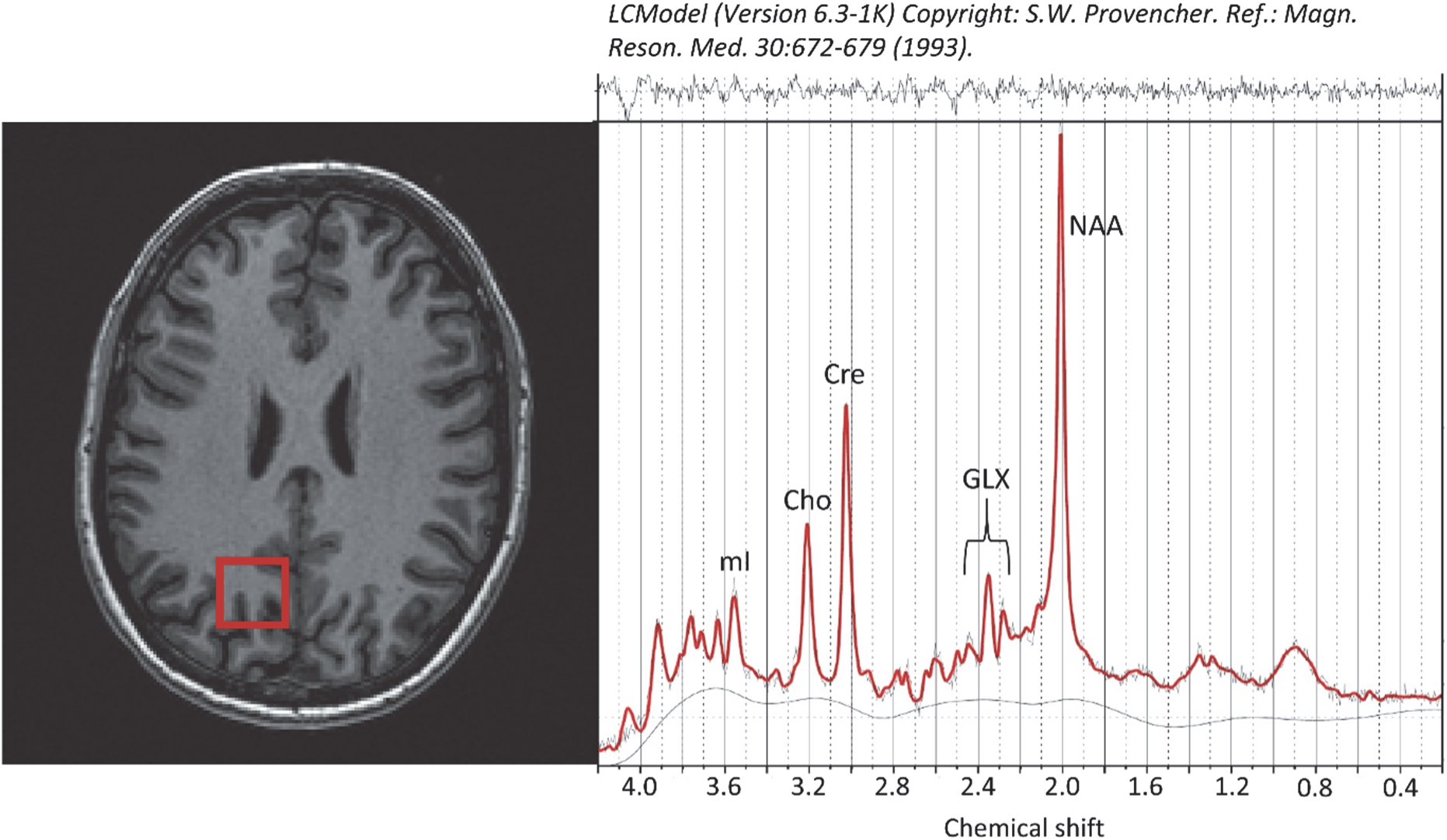
Voxel position in the left parietal lobe (red rectangle on axial T1 image) and representative magnetic resononance spectrum. mI = myo-inositol, Cho = Cholin, Cre = Creatine, GLX = glutamate + glutamine, NAA = N-acetylaspartate

### Statistics

Changes in metabolite concentrations and reduction in SWA were evaluated using paired samples t-tests. To correct for multiple comparisons, a false discovery rate (FDR) correction was applied. In order to correct for individual differences in absolute power values, the decrease in SWA was calculated as the percentage reduction from the NREM sleep episode with maximum SWA (= 100 %) to the last NREM sleep episode. Overnight changes in metabolites concentrations were calculated in the same way, with evening levels set to 100 % and overnight changes calculated as percentage change. Additionally, changes in EEG power were calculated for 4 additional classical frequency bands (5 - 8 Hz, 8 - 10 Hz, 10 - 12 Hz, 12 - 15 Hz) in the same cycles as selected for SWA.

In order to assess the association between changes in GLX levels and SWA, Pearson correlation coefficients between the decrease in SWA at every electrode and the overnight changes in GLX levels were calculated. To correct for multiple comparisons, statistical nonparametric mapping (SnPM) using a suprathreshold cluster analysis was applied (Huber et al., 2004, Nichols and Holmes, 2001). To assess whether the overnight change in GLX is specifically associated with SWA, the percentage change in GLX was correlated to the percentage change in the 4 additional frequency bands (5 - 8 Hz, 8 - 10 Hz, 10 - 12 Hz, 12 - 15 Hz). All analyses were performed with the software package MATLAB (MathWorks). The significance level was set at *p* < 0.05 (two-tailed).

## Results

We obtained good quality MR spectra from all subjects at both time points (Fig. 1 for an example). In a first step, we compared evening and morning levels of GLX and 5 other metabolites with robust peak detection [N-acetylaspartate (NAA), N-acetyl aspartate + N-acetyl aspartylglutamate (NAA + NAAG), glycerophosphocholine + phosphocholine (GPC + PCh), myo-Inositol (mI) and creatine (Cre)]. Significant overnight changes were only observed for GLX / H_2_O, where 13 out of 16 subjects showed an overnight reduction (-6.63 ± 1.59 %, *p* = 0.006, FDR corrected, effect size = 1.0; Fig. 2 & Tab. 1). No significant changes were observed in any other metabolite (all *p* > 0.4, effect size ≤ 0.3). Overnight changes in levels of metabolites were not driven by variations in voxel composition, since grey, white and CSF content of the evening and morning voxels showed no differences (all *p* > 0.6). In a next step, we tested whether metabolite concentrations were related to sleep stages. We found no significant correlations between any of the sleep stages and metabolite concentrations in the evening, the morning or the overnight change (all *p* > 0.4, FDR corrected; Tab. 2). Because the gradual decrease of SWA over the night is thought to reflect the restorative function of sleep, we tested for an association between the decrease in SWA and the overnight reduction in GLX. First, we confirmed that SWA displayed a highly significant decrease in all 109 electrodes from the maximal to last NREM sleep episode (mean reduction of 70.1 ± 3.1 %, *p* < 0.001, FDR corrected; Fig. 3). Electrode-wise Pearson correlations between the decrease in SWA and the overnight reduction in GLX revealed a significant cluster of 69 electrodes (Fig. 4A). The mean correlation within this cluster was *r* = 0.57, *p* = 0.02 (Fig. 4B). There was no significant correclation between all night SWA (averaged across all electrodes) and changes in GLX (*p* = 0.6). We then calculated the percentage change of EEG power in the 4 other frequency bands within this cluster of 69 electrodes and correlated it with the reduction in GLX. Results revealed that besides SWA, there was a significant association between GLX and power in the theta frequency band (5 - 8 Hz; *r* = 0.53, *p* = 0.03; all other bands *p* > 0.4; Fig. 4C).

**Figure 2.**
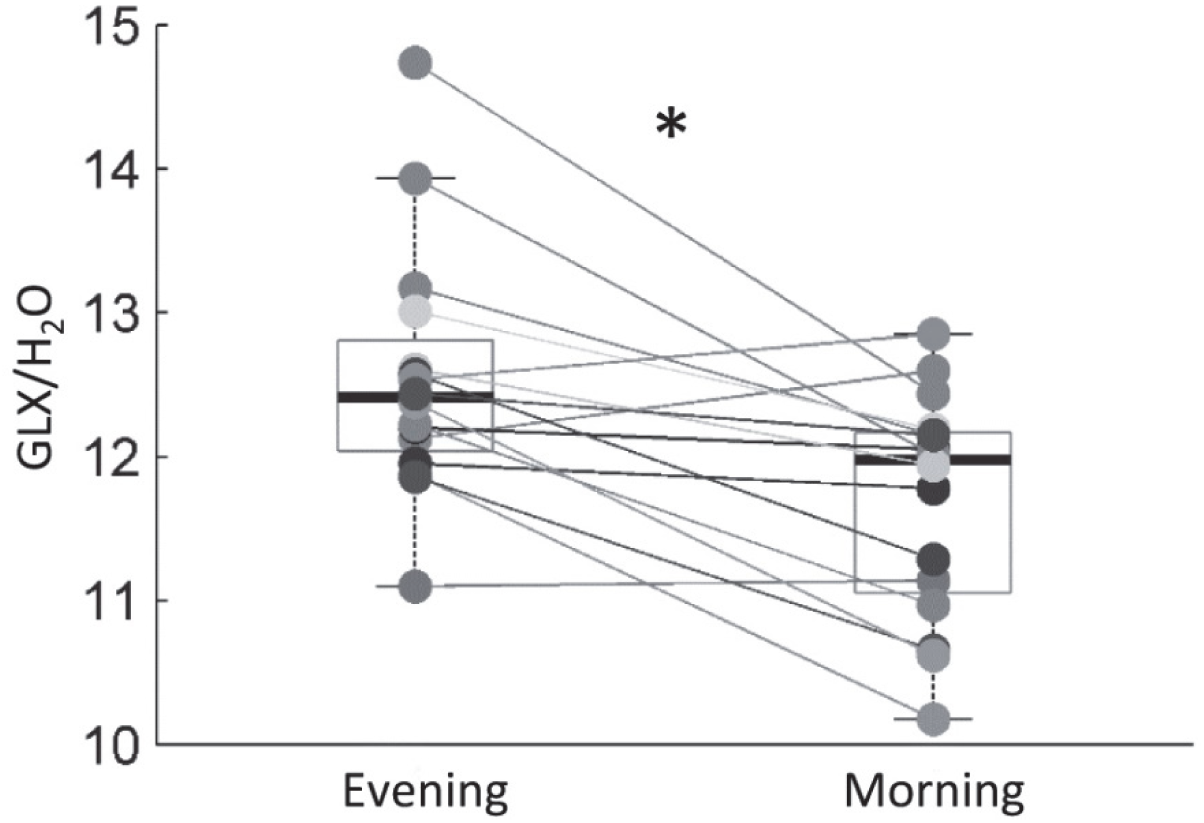
Changes in water-scaled glutamate + glutamine (GLX) levels from evening to morning. Mean (± s.e.m.) reduction from evening to morning was 6.63 (± 1.59 %, * *p* = 0.006, FDR corrected).

**Table 1.** Water-scaled metabolite concentrations in the evening and the morning and their change across the night.

| (n=16) | Evening |  | Morning |  | Overnight change |  |
| --- | --- | --- | --- | --- | --- | --- |
|  | mean | s.e.m. | mean | s.e.m. | (%) | s.e.m. |
| GLX | 12.6 | ± 0.22 | 11.7 | ± 0.20 | -6.6* | ± 1.59 |
| NAA | 11.6 | ± 0.13 | 11.5 | ± 0.12 | -0.3 | ± 1.21 |
| NAA+NAAG | 12.6 | ± 0.15 | 12.5 | ± 0.12 | -0.9 | ± 1.51 |
| GPC+PCh | 1.6 | ± 0.05 | 1.6 | ± 0.06 | -2.0 | ± 1.55 |
| mI | 3.7 | ± 0.08 | 3.6 | ± 0.11 | -3.2 | ± 2.46 |
| Cre | 7.5 | ± 0.14 | 7.5 | ± 0.14 | -0.7 | ± 1.21 |

GLX = glutamate + glutamine, NAA = N-acetyl aspartate, NAA + NAAG = N-acetyl aspartate + N-acetyl aspartylglutamate, GPC + PCh = glycerophosphocholine + phosphocholine, mI = myo-Inositol, Cre = Creatine
\* $p < 0.05$ , FDR corrected

**Table 2.** Sleep architecture.

| (n=16) | mean | s.e.m. |
| --- | --- | --- |
| Time in bed (min) | 458.6 | ±6.30 |
| Sleep latency (min) | 17.1 | ± 2.68 |
| Total sleep time (min) | 421.4 | ± 7.09 |
| Sleep efficiency (%) | 91.9 | ± 1.14 |
| Wake after sleep onset (min) | 24.0 | ± 3.51 |
| NREM sleep (min) | 321.1 | ± 5.15 |
| N1 (min) | 25.8 | ± 2.96 |
| N2 (min) | 207.8 | ± 8.71 |
| N3 (min) | 87.6 | ± 8.61 |
| REM sleep (min) | 100.3 | ± 6.03 |

NREM sleep = non-rapid eye movement sleep, REM sleep = rapid eye movement sleep
Sleep efficiency = time in bed divided by total sleep time (NREM+REM)

**Figure 3.**
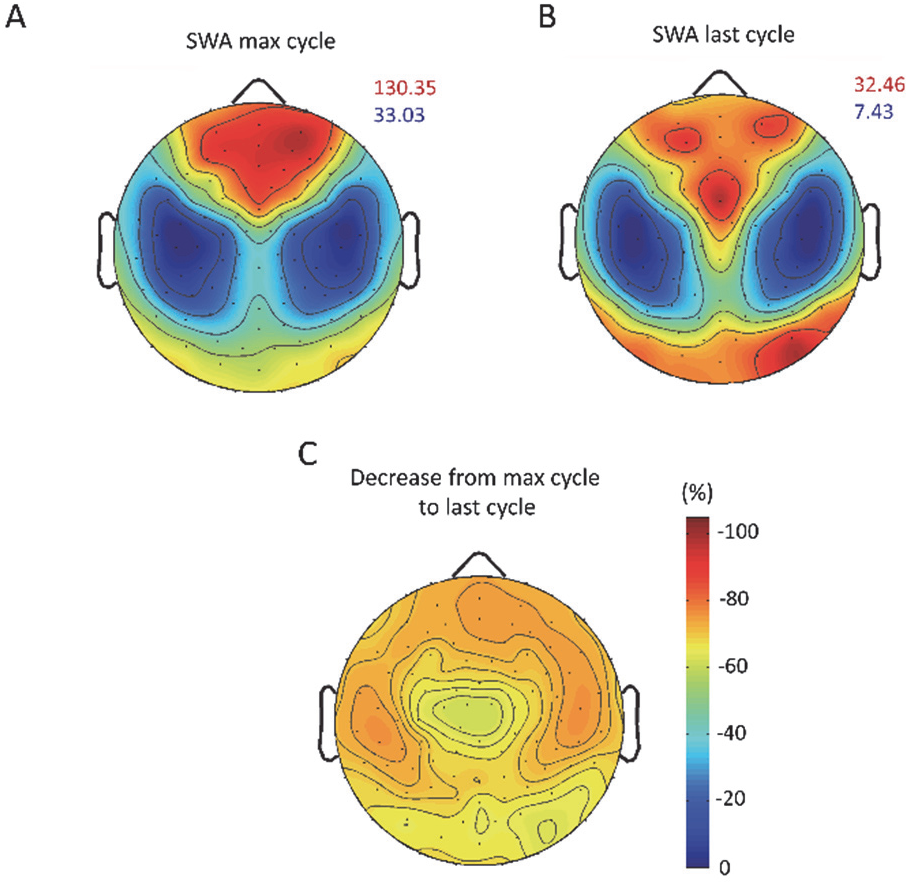
Topographical distribution of EEG slow wave activity (SWA). (A) Topographical map of SWA during the NREM sleep episode with maximum SWA and (B) during the last NREM sleep episode, respectively scaled to their maximum (red) and minimum (blue) power values (µV2 / Hz). (C) Electrode wise reduction in SWA from NREM sleep episode with maximal SWA to last NREM sleep episode (%), minimal reduction −62 %, maximal reduction −76 %. Reduction was significant at all electrodes (all *p* < 0.001, FDR corrected).

**Figure 4.**
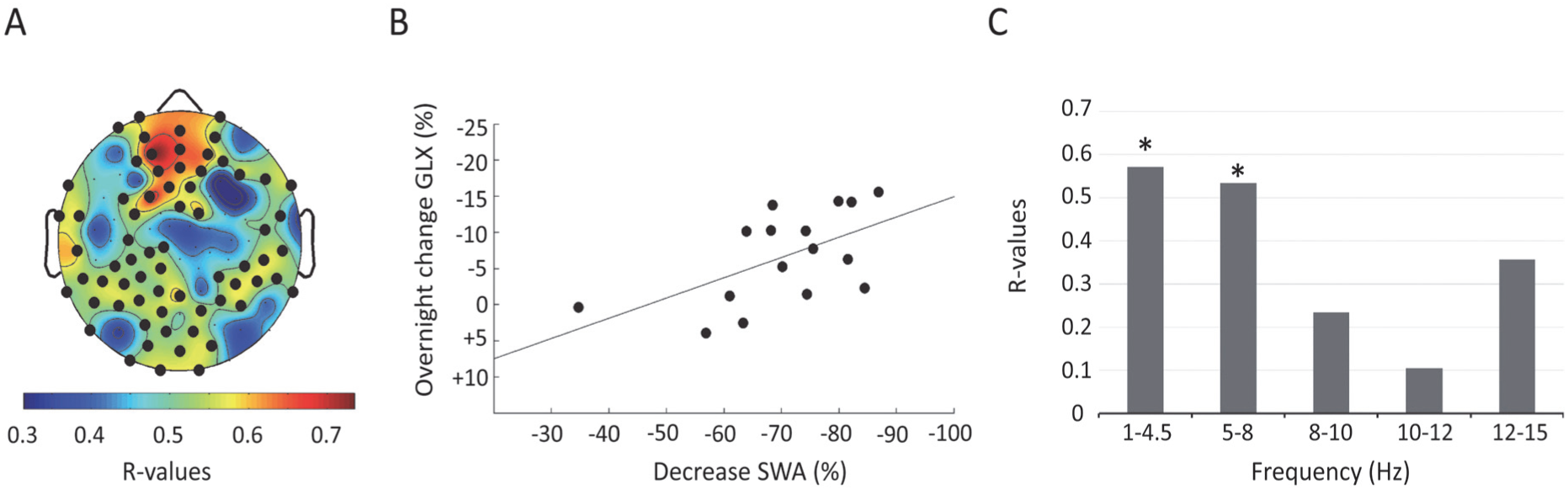
Correlation between the decrease in SWA and overnight changes in GLX. (A) Results of the electrode-wise Pearson correlations between the decrease of SWA (%) and the overnight change in GLX (%) plotted on the planar projection of the hemispheric scalp model. Bold black dots indicate electrodes showing significant correlations [defined by statistical nonparametric mapping (SnPM); suprathreshold cluster analysis to control for multiple comparison (Nichols & Holmes, 2002)]. (B) Association between the average decrease of SWA (%) in the significant cluster electrodes and overnight changes in GLX (%), *r* = 0.57, *p* = 0.02. (C) Pearson correlation coefficients of changes in GLX (%) and changes (%) in power in the 4 additional frequency bands. * *p* < 0.05

## Discussion

In this study, we assessed whether overnight changes in levels of GLX are measureable by ^1^H-MRS in healthy young adults. Indeed, we found a highly significant overnight reduction in GLX, whereas other metabolites did not display an overnight change. This observation suggests that our findings are specific to GLX and do not arise from changes in the unsuppressed water signal used for scaling of the metabolite values. Next, we tested if changes in GLX are correlated to EEG SWA, our primary electrophysiological marker of the restorative function of sleep. We found a positive correlation between the decrease in GLX and SWA in the course of the night. Both of these results fit to studies in rats demonstrating changes in the level of glutamate as a function of sleep and wake. More specifically, it has been shown that levels of glutamate decrease during NREM sleep and that this decrease is positively correlated with SWA (Dash et al., 2009). An obvious limitation of our study is that we were not able to robustly differentiate between glutamine and glutamate for technical reasons. Scanning at stronger magnetic fields (e.g. 7 Tesla) might overcome this limitation. However, even with improved differentiation, ^1^H-MRS does not allow to separate between intracellular and extracellular glutamate.

Our experimental design allowed an MRS scan of a single voxel. We therefore cannot draw any conclusions for the whole brain. Interestingly, although we measured GLX levels in this single voxel sized 20 x 20 x 20 mm^3^ in the left parietal lobe, we found a rather global correlation with SWA over a widespread cluster of electrodes. Within this cluster, correlation was strongest in the frontal region followed by correlations in the parietal and occipital lobes. One explanation for this rather global effect might be that glutamate, the most abundant excitatory neurotransmitter, is highly widespread in the brain (Huganir and Nicoll, 2013, Lee and Kirkwood, 2011) and its release strongly depends on overall neuromodulation and neuronal firing (Tzingounis and Wadiche, 2007). Moreover, as shown in Figure 3, SWA decreases very consistently across the entire cortex, which may contribute to a rather global relationship with GLX. To further investigate a region specific relationship, future studies should quantify glutamate in additional brain areas.

A critical limitation of our study is that GLX was assessed at two time points differing in circadian phase. As circadian rhythms exert strong influences at various levels, ranging from cognitive activity (Waterhouse, 2010) to the level of neuromodulation (Stanley et al., 1989), our difference in GLX could be explained by a difference in circadian time. However, the specific correlation with SWA speaks against such an explanation because the recovery function of sleep, as reflected by SWA, is to a large extent independent of circadian rhythms (Borbély and Achermann, 2005). Future studies incorporating sleep deprived participants may be able to confirm whether the apparent decrease in GLX in the morning is due to sleep or the circadian time. The correlation between the changes in GLX extended into power in the theta frequency range (5 - 8 Hz), albeit it was less strong than for SWA. This is not surprising, since SWA and power in the theta range seem to be regulated together during sleep, as prolonged wakefulness not only results in an increase of SWA but also extends into higher frequencies including the theta frequency band (Finelli et al., 2000).

Since our study is purely correlative, future studies need to prove causality. Moreover, as done in animal studies, it would be critical to see the continuous changes in glutamate in the course of sleep. Such studies could investigate whether or not our observed reduction in GLX is related to deep sleep’s ability to reduce high energy demands and/or down-regulation of synaptic strength, preventing saturation of neuronal networks and ensuring efficient cortical functioning. If indeed sleep SWA plays a critical role in glutamate homeostasis, MRS measured GLX levels might be a promising biomarker of the recuperative function of sleep.

## References

Bollmann S, Ghisleni C, Poil SS, Martin E, Ball J, Eich-Hochli D, Edden RA, Klaver P, Michels L, Brandeis D & O’Gorman RL (2015) Developmental changes in gamma-aminobutyric acid levels in attention-deficit/hyperactivity disorder. Transl Psychiatry 5, e589.

Borbély AA & Achermann P (2005) Sleep homeostasis and models of sleep regulation. In: Principles and Practice of Sleep Medicine, (eds.) M. H. Kryger, T. Roth & W. C. Dement, pp. 405–417. Philadelphia: Elsevier Saunders.

Dash MB, Douglas CL, Vyazovskiy VV, Cirelli C & Tononi G (2009) Long-term homeostasis of extracellular glutamate in the rat cerebral cortex across sleep and waking states. J Neurosci 29, 620–629.

Delorme A & Makeig S (2004) EEGLAB: an open source toolbox for analysis of single-trial EEG dynamics including independent component analysis. J Neurosci Methods 134, 9–21.

Dlabac-de Lange JJ, Liemburg EJ, Bais L, van de Poel-Mustafayeva AT, de Lange-de Klerk ES, Knegtering H & Aleman A (2017) Effect of Bilateral Prefrontal rTMS on Left Prefrontal NAA and Glx Levels in Schizophrenia Patients with Predominant Negative Symptoms: An Exploratory Study. Brain Stimul 10, 59–64.

Feinberg I & Floyd TC (1979) Systematic trends across the night in human sleep cycles. Psychophysiology 16, 283–291.

Finelli LA, Baumann H, Borbély AA & Achermann P (2000) Dual electroencephalogram markers of human sleep homeostasis: correlation between theta activity in waking and slow-wave activity in sleep. Neuroscience 101, 523–529.

Hanlon EC, Faraguna U, Vyazovskiy VV, Tononi G & Cirelli C (2009) Effects of skilled training on sleep slow wave activity and cortical gene expression in the rat. Sleep 32, 719–729.

Huang Z, Davis HI, Yue Q, Wiebking C, Duncan NW, Zhang J, Wagner NF, Wolff A & Northoff G (2015) Increase in glutamate/glutamine concentration in the medial prefrontal cortex during mental imagery: A combined functional mrs and fMRI study. Hum Brain Mapp 36, 3204–3212.

Huber R, Ghilardi MF, Massimini M, Ferrarelli F, Riedner BA, Peterson MJ & Tononi G (2006) Arm immobilization causes cortical plastic changes and locally decreases sleep slow wave activity. Nat Neurosci 9, 1169–1176.

Huber R, Ghilardi MF, Massimini M & Tononi G (2004) Local sleep and learning. Nature 430, 78–81.

Huganir RL & Nicoll RA (2013) AMPARs and synaptic plasticity: the last 25 years. Neuron 80, 704–717.

Iber C, Ancoli-Israel S, Chesson A & Quan SF (2007) The AASM manual for the scoring of sleep and associated events: rules, terminology and technical specifications. (ed.) A. A. o. S. Medicine, 1st edition. Westchester, Illinois.

Krause AJ, Simon EB, Mander BA, Greer SM, Saletin JM, Goldstein-Piekarski AN & Walker MP (2017) The sleep-deprived human brain. Nat Rev Neurosci 18, 404–418.

Lee HK & Kirkwood A (2011) AMPA receptor regulation during synaptic plasticity in hippocampus and neocortex. Semin Cell Dev Biol 22, 514–520.

Nichols TE & Holmes AP (2001) Nonparametric permutation tests for functional neuroimaging: a primer with examples. Hum. Brain Mapp. 15, 1–25.

Robertson, D. M. W., Van Amelsvoort, T., Daly, E., Simmons, A., Whitehead, M., Morris, R. G., … & Murphy, D. G. M. (2001). Effects of estrogen replacement therapy on human brain aging: an in vivo 1H MRS study. Neurology, 57(11), 2114–2117.

Rowland LM, Kontson K, West J, Edden RA, Zhu H, Wijtenburg SA, Holcomb HH & Barker PB (2013) In vivo measurements of glutamate, GABA, and NAAG in schizophrenia. Schizophr Bull 39, 1096–1104.

Stanley BG, Schwartz DH, Hernandez L, Hoebel BG & Leibowitz SF (1989) Patterns of extracellular norepinephrine in the paraventricular hypothalamus: relationship to circadian rhythm and deprivation-induced eating behavior. Life Sci 45, 275–282.

Tononi G & Cirelli C (2014) Sleep and the price of plasticity: from synaptic and cellular homeostasis to memory consolidation and integration. Neuron 81, 12–34.

Turrigiano GG (2008) The self-tuning neuron: synaptic scaling of excitatory synapses. Cell 135, 422–435.

Tzingounis AV & Wadiche JI (2007) Glutamate transporters: confining runaway excitation by shaping synaptic transmission. Nat Rev Neurosci 8, 935–947.

Van Dongen HP, Maislin G, Mullington JM & Dinges DF (2003) The cumulative cost of additional wakefulness: dose-response effects on neurobehavioral functions and sleep physiology from chronic sleep restriction and total sleep deprivation. Sleep 26, 117–126.

Vyazovskiy VV, Cirelli C, Pfister-Genskow M, Faraguna U & Tononi G (2008) Molecular and electrophysiological evidence for net synaptic potentiation in wake and depression in sleep. Nat Neurosci 11, 200–208.

Vyazovskiy VV & Harris KD (2013) Sleep and the single neuron: the role of global slow oscillations in individual cell rest. Nat Rev Neurosci 14, 443–451.

Vyazovskiy VV, Olcese U, Lazimy YM, Faraguna U, Esser SK, Williams JC, Cirelli C & Tononi G (2009) Cortical firing and sleep homeostasis. Neuron 63, 865–878.

Waterhouse J (2010) Circadian rhythms and cognition. Prog Brain Res 185, 131–153.

Xie L, Kang H, Xu Q, Chen MJ, Liao Y, Thiyagarajan M, O’Donnell J, Christensen DJ, Nicholson C, Iliff JJ, Takano T, Deane R & Nedergaard M (2013) Sleep drives metabolite clearance from the adult brain. Science 342, 373–377.

